# Inhibition of oxytocin neurons during key periods of development has long-term behavioral and body composition effects

**DOI:** 10.1101/2025.11.13.688271

**Authors:** Fabienne Schaller, Marie-Sophie Alifrangis, Pierre-Yves Barelle, Catarina Santos, Roman Tyzio, Alessandra Bertoni, Felix Omnes, Yasmine Belaidouni, Francesca Bader, Emilie Pallesi-Pocachard, Jean-Luc Gaiarsa, Sebastien G Bouret, Françoise Muscatelli

**Affiliations:** Institut de Neurobiologie de la Méditerranée (INMED), INSERM, Aix Marseille Université, Marseille, France; Univ. Lille, INSERM, CHU Lille, Laboratory of Development and Plasticity of the Neuroendocrine Brain, Lille Neuroscience and Cognition, Lille, France; Phenotype Expertise, Marseille, France

## Abstract

**Background:** Oxytocin (OT) is a key neuromodulator of social behavior in mammals, and accumulating evidence supports the existence of a critical period for OT action during infancy. However, other developmental windows remain poorly explored, and it remains unclear whether OT exert distinct functions depending on the timing of its activity. In this study, we aimed to determine whether specific developmental stages exist during which OT-expressing neurons play a decisive role with long-term consequences.

**Methods:** We used a chemogenetic approach to transiently inhibit OT-expressing neurons during three postnatal developmental periods: infancy, the juvenile period, and young adulthood in male and female mice. Behavioral and metabolic outcomes were then assessed longitudinally. We also examined the effects of inhibiting OT neurons during late fetal stages and at birth.

**Results:** Social memory was consistently impaired in males, regardless of the timing of neuronal inactivation. The most pronounced behavioral effects were observed following inhibition during infancy in both sexes. Metabolically, adult males from all cohorts exhibited increased body weight, whereas increased fat mass and adipocyte size hypertrophy were specifically observed following inhibition during the juvenile period. Notably, inhibition of OT-expressing neurons around the time of birth resulted in delayed parturition and altered neonatal feeding behavior.

**Limitations:** OT-expressing neurons release multiple signaling molecules. However, converging evidence suggests that the observed phenotypes are primarily attributable to OT deficiency. Although this study demonstrates long-term behavioral and metabolic consequences of transient OT neuron inhibition, further experiments are required to elucidate underlying mechanisms and identify additional effects.

**Conclusions:** Transient inhibition of OT-expressing neurons during distinct postnatal developmental periods leads to long-lasting effects on social behavior and metabolism, with outcomes depending on both the timing of inhibition and sex. Furthermore, our findings reveal an unexpected role for fetal/neonatal OT neurons in regulating the timing of birth and early feeding behavior.

## INTRODUCTION

Oxytocin (OT) is involved in numerous physiological processes and acts as a powerful neuromodulator of social behavior in primates and rodents [1, 2]. Over the last two decades, growing evidence has highlighted the role of OT in neurodevelopment [3, 4]. In particular, postnatal OT manipulations have been well studied in prairie voles, where their effects depend on sex, dose, and brain region [5, 6].

OT has been implicated in the pathophysiology of several neurodevelopmental disorders, including Autism Spectrum Disorders (ASD), Prader–Willi syndrome (PWS), and Schaaf–Yang syndrome [7, 8], as well as in stress and anxiety-related disorders [9] and post-traumatic stress disorder (PTSD) [10]. Accordingly, OT-based treatments have been administered to patients with ASD and PWS [11], stress-related psychiatric disorders, and PTSD [12]. Although some beneficial effects have been reported, outcomes remain inconsistent. This variability likely reflects heterogeneity in patient population (*e.g*., age, aetiology, and environment), as well as differences in administration methods, including dose, duration, and delivery methods that are not yet grounded in robust scientific evidence [11, 12, 13, 14]. However, OT treatment in infants [15, 16] and in children with PWS [17, 18, 19, 20] has shown more consistent and long-lasting improvements in feeding and social behaviors [16].

In several rodent models, including knockouts (KO) of ASD risk genes (*Magel2, Cntnap2, Fmr1, Nlgn3, Shank3, Oprm1, Pten*), BTBR mice, and valproic acid-exposed rodents, the OT system has been implicated in social deficits [21]. Moreover, OT signaling represents a convergent pathway in autism [22, 23]. Importantly, early-life OT treatment restores social behavior in many of these models, with effects persisting into adulthood. This has been demonstrated in *Magel2* KO [24, 25], *Cntnap2* KO [26], *Fmr1* KO [27], the 22q11.2 hemideletion syndrome mouse model [28], valproic acid-treated mice [29], and a rat model of PTSD [30].

Together, these findings support the concept of an early-life OT-mediated hormonal imprinting that shapes neural circuits involved in social behavior over the long term, suggesting the existence of a critical developmental window during infancy. Consistent with this idea, recent work indicates that the early postnatal period may represent a unique critical window during which OT promotes prosocial behaviors by regulating the GABA shift [29], as we previously reported [25]. However, additional sensitive periods during which OT regulates specific behavioral or metabolic processes may also exist. For example, social conditioned place preference in mice is influenced by OT signaling during a developmental window spaning from weaning to early adulthood, but not during infancy [31].

Despite these advances, no study has directly examined the effects of transient inhibition of OT neurons across distinct developmental periods. Such an approach is essential for understanding the role of OT signaling in the pathogenesis of neurodevelopmental disorders, and for developing therapeutic strategies aimed at targeting, or potentially reopening, these sensitive periods [31]. To address this gap, we used chemogenetics tools to inhibit OT-expressing neurons at three developmental stages: the first postnatal week, the juvenile (pubertal) period in male and female mice, and young adulthood (P60). We assessed both behavioral and metabolic outcomes. In addition, to evaluate the importance of OT produced by newborns, OT neurons were inhibited from embryonic day 18 until birth. These time points were selected to capture key stages of brain development, including the maturation and connectivity of OT neurons [32] and the dynamic expression of OT receptors [33].

## MATERIALS ET METHODS

### Study approval

All experiments were performed in accordance with the Guide for the Care and Use of Laboratory Animals (N.R.C., 1996) and the European Communities Council Directive of September 22th 2010 (2010/63/EU, 74) and the approved protocol (APAFIS#21251-2019092314503551 for the studies performed in Marseille and APAFIS# 13387–2017122712209790 for the studies performed in Lille) by the Ethical Committee of the French Ministry of Agriculture. Animals were group-housed with a 12-h light-dark cycle (light from 7pm to 7 am and dark from 7am to 7pm), at a constant temperature (22 ± 1°C) with *ad libitum* access to standard chow and water.

### Sex as a biological variable

Our study examined male and female animals. Both sexes were pooled when no sex differences were found.

| Species and genetic background | Designation | Reference | Additional informations |
| --- | --- | --- | --- |
| Mice, C57BL6/J | WT |  | Charles River |
| Mice, C57BL6/J | OT::Cre | (Wu *et al.*) [34] | JAX stock #024234 |
| Mice, C57BL6/J | Ai14(LSL-tdTomato) | (Madisen *et al.*) [35] | JAX Stock 007914 |
| Mice, C57BL6/J | R26-LSL-hM4Di-DREADD | (Zhu *et al.*)[36] | JAX stock #026219 |

### Mouse strains

The R26-LSL-hM4Di-DREADD mice were on a C57Bl6/N genetic background, which is not suitable for conducting behavioral analyses, due to a mutation in the gene encoding Cytoplasmic FMR interacting protein 2, which regulates behavior [37]. We performed over 8 back crosses with C57BL6/J mice and validated by sequencing the deletion of the Cyfip2 mutation in mice used in the breedings to generate all the experimented mice.

### Chemogenetic inactivation by intra-peritoneal or oral administration of Compound 21

Chemogenetic experiments were performed on OT::Cre-/+;R26-LSL-hM4Di-DREADD/R26-LSL-hM4Di-DREADD (named Cre+) male and female mice. A subcutaneaous (in infancy) or intraperitoneal (at adolescence or adulthood) injection of C21 (DREADD agonist 21 dihydrochloride, Hello Bio, Cat. No. HB HB6124 Hello Bio Ltd, Ireland) at 2.5mg/kg (diluted in NaCl 0.9%) was performed twice a day (one at 9am and one at 5pm). To inactivate OT-expressing neurons in late embryos and newborns, we added C21(0.25 mg/mL) in drinking water of the female from Gestational Day 18 to one day after birth. Gestational female at this stage have a body weight around 40g and consume ∼5 ml of water per day [38]. Accordingly, C21 dissolved in drinking water at 0.25mg/ml results in 4-6mg/kg of C21 active dose at any time since a mouse drinks all over the 24 hours and the duration of action of a single dose of C21 is around 4-6hours [39]. Mouse cage water containing C21 and was carefully monitored and changed for fresh solution every day.

### Immunohistochemistery

P20 mice were deeply anaesthetized with intraperitoneal injection of the ketamine/xylazine mixture and transcardially perfused with 0.9% NaCl saline followed by Antigenfix (Diapath, cat #P0014). Brains were post-fixed in Antigenfix overnight at 4°C and included in agar 4%. 50 μm-thick coronal sections were sliced using a vibratom (Zeiss) and stored in PBS at 4°C. Floating slices (of the hippocampal region corresponding to slices 68 to 78 on Allan Brain Atlas) were incubated for 1 hour with blocking solution containing 0.1% (v/v) Triton X-100, 10% (v/v) normal goat serum (NGS) in PBS, at room temperature. Sections were then incubated with primary antibodies diluted in incubation solution (0.1% (v/v) Triton X-100, 3% (v/v) NGS, in PBS), overnight at 4C°. After 3 x 10 min washes in PBS, brain sections were incubated with secondary antibodies diluted in the incubation solution, for 2 hours at RT. Sections were washed 3 x 10 min in PBS and mounted in Flouromount-G (EMS, cat #17984-25).

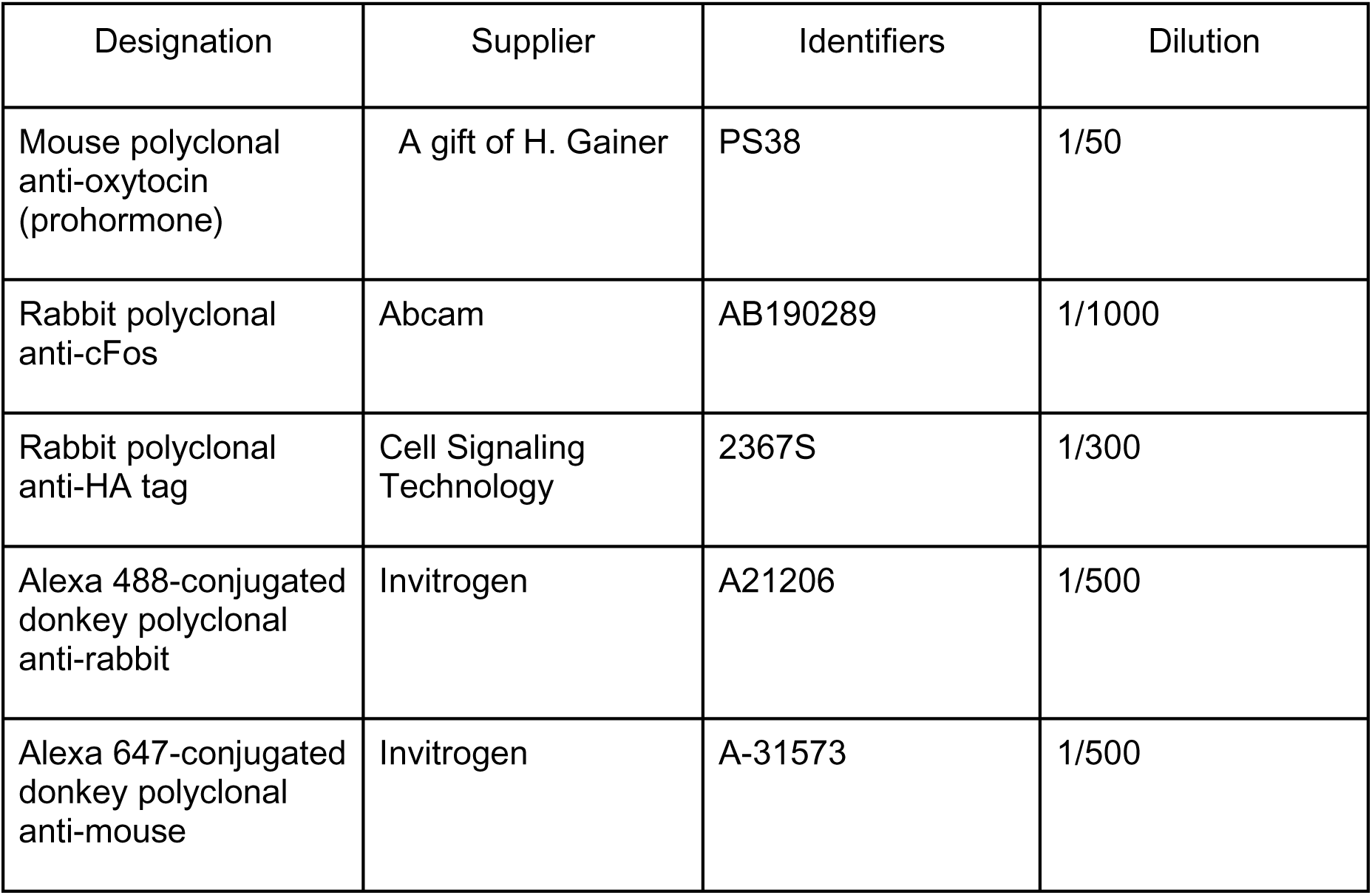

Images were captured using a fluorescence microscope (Zeiss Axioplan 2 microscope with an Apotome module).

### Electrophysiological experiments

#### PVH slice preparations and recording of OT neurons

OT-Cre::Ai14(LSL-tdTomato)::R26-LSL-hM4Di-DREADD mice at P5 were anesthetized using 5% isoflurane and rapidly decapitated. Coronal slices containing the PVH (350 µm) were generated using a vibratome (VT 1200S, Leica, Germany). Slices were allowed to equilibrate before use for 1h in standard recording aCSF (in mM: 124 NaCl, 3 KCl, 1.2 NaH2PO4, 1.2 MgSO4, 25 NaHCO3, 11 D-glucose, and 2 CaCl2, saturated with 95% O2/5% CO2, pH 7.4, ∼300 mos M) at room temperature. During the experiment the recording chamber was superfused at ∼3 mL/min with the standard recording aCSF.

Electrodes (resistance ∼3.0-6.0 MOhm) were prepared using a Narishige micropipette puller (PC-100) and filled with (in mM): 130 K^+^-gluconate, 10 NaCl, 1.1 EGTA, 1 CaCl2, 10 HEPES, 1 MgCl2, 2 Mg-ATP, 0.2 Na-GTP, pH 7.3, and ∼280 mosM. Recording pipettes were guided using a piezoelectric manipulator (Luigs & Neumann, GmbH). PVH OT-expressing neurons expressing tdTomato were identified in Cre^+^ mice using a Hamamatsu scientific camera. In Cre− mice, unlabeled neurons in caudal PVH slices were recorded. Following initial membrane rupture the resting membrane potential was measured. Resting membrane potential, spontaneous action potentials, and evoked action potentials were recorded in current clamp mode. The DREADD agonist C21 (10 µM) was applied for 5 min. Data obtained from PVH slice experiments were acquired in pClamp 10.7 using a HEKA EPC 10 Double amplifier (HEKA Elektronik), filtered at 2 kHz and sampled at 20 kHz. Data were analyzed using Clampfit 10.7 and MiniAnalysis (Synaptosoft, USA).

#### Hippocampal slice preparation and recordings of pyramidal CA3 neurons

Brains were removed and immersed into ice-cold (2-4°C) artificial cerebrospinal fluid (ACSF) with the following composition (in mM): 126 NaCl, 3.5 KCl, 2 CaCl_2_, 1.3 MgCl_2_, 1.2 NaH_2_PO_4_, 25 NaHCO_3_ and 11 glucose, pH 7.4 equilibrated with 95% O_2_ and 5% CO_2_. Hippocampal slices (400 µm thick) were cut with a vibratome (Leica VT 1000s, Germany) in ice cold oxygenated choline-replaced ACSF and were allowed to recover at least 90 min in ACSF at room (25°C) temperature. Slices were then transferred to a submerged recording chamber perfused with oxygenated (95% O_2_ and 5% CO_2_) ACSF (3 ml/min) at 34°C.

#### Whole-cell patch clamp recordings

Were performed from P20-P25 CA3 pyramidal neurons in voltage-clamp mode using an Axopatch 200B (Axon Instrument, USA). To record the spontaneous and miniature synaptic activity, the glass recording electrodes (4-7 M) were filled with a solution containing (in mM): 100 KGluconate, 13 KCl, 10 HEPES, 1.1 EGTA, 0.1 CaCl_2_, 4 MgATP and 0.3 NaGTP. The pH of the intracellular solution was adjusted to 7.2 and the osmolality to 280 mOsmol l^-1^. The access resistance ranged between 15 to 30 M. With this solution, the GABA_A_ receptor-mediated postsynaptic current (GABAA-PSCs) reversed at - 70mV. GABA-PSCs and glutamate mediated synaptic current (Glut-PSCs) were recorded at a holding potential of -45mV. At this potential GABA-PSC are outwards and Glut-PSCs are inwards. Spontaneous synaptic activity was recorded in control ACSF and miniature synaptic activity was recorded in ACSF supplemented with tetrodotoxin (TTX, 1µM). Spontaneous and miniature GABA-PSCs and Glut-PSCs were recorded with Axoscope software version 8.1 (Axon Instruments) and analyzed offline with Mini Analysis Program version 6.0 (Synaptosoft).

### Behavioral studies and Metabolic studies

See Supplementary Information

### Statistical Analysis

Statistical analyses were performed using GraphPad Prism (GraphPad Software, Prism 7.0 software, Inc, La Jolla, CA, USA). All statistical tests were two-tailed and the level of significance was set at P<0.05. Appropriate tests were conducted depending on the experiment; tests are indicated in the figure legends or detailed in supplementary statistical file. Values are indicated as Q2 (Q1, Q3), where Q2 is the median, Q1 is the first quartile and Q3 is the third quartile when non-parametric tests were performed and scatter dot plots report Q2 (Q1,Q3) or as mean ± SEM when parametric tests were performed usually in histograms. N refers to the number of animals, while n refers to the number of brain sections or hippocampi or cells recorded.

Mann-Whitney (MW) non-parametric test or t-test (parametric test) were performed to compare two matched or unmatched groups. ANOVA or Kruskal-Wallis tests were performed when the different groups have been experienced in the same experimental design only; if this was not the case, MW or t-test were used. One-way ANOVA followed by Bonferroni or Dunnett’s or Tukey’s post-hoc tests were used to compare three or more independent groups. Two-way ANOVA followed by Bonferroni post-hoc test was performed to compare the effect of two factors on unmatched groups. *: p< 0.05; **: p <0.01; ***: p<0.001; ****: p<0.0001.

## RESULTS

### Chemogenetic inhibition of OT-expressing neurons and experimental design

To transiently inhibit OT neurons, we generated a transgenic mouse model (OT-Cre::hM4Di-HA) expressing the inhibitory Gαi/o-coupled receptor hM4Di (R26-LSL-hM4Di-HA) to selectively in OT-expressing neurons using the Designer Receptors Exclusively Activated by Designer Drugs (DREADD) system. Specific breeding strategies yielded OT-Cre heterozygous::hM4Di-HA homozygous mice (hereafter Cre+) and hM4Di-HA homozygous littermates lacking Cre (Cre−, controls) (Fig. 1a). Selective expression of hM4Di-HA in OT neurons was validated (Fig. 1b). Electrophysiological recordings in brain slices from P3-P5 pups demonstrated that the DREADD agonist Compound 21 (C21) effectively suppressed action potential firing in OT neurons from Cre+ mice, but not Cre− controls (Fig. 1c). C21 was selected for its high potency, selectivity, ∼6 hour-duration of action, good brain penetrance [40, 41] and minimal off-target behavioral effects [41, 42]. Importantly, prior validation confirmed that transient C21 treatment, performed at the threedevelopmental stages studied, does not alter the integrity of OT-expressing neurons at adulthood [43]. Following transient inhibition, mice underwent a battery of behavioral tests (Fig. 1d).

**Figure 1.**
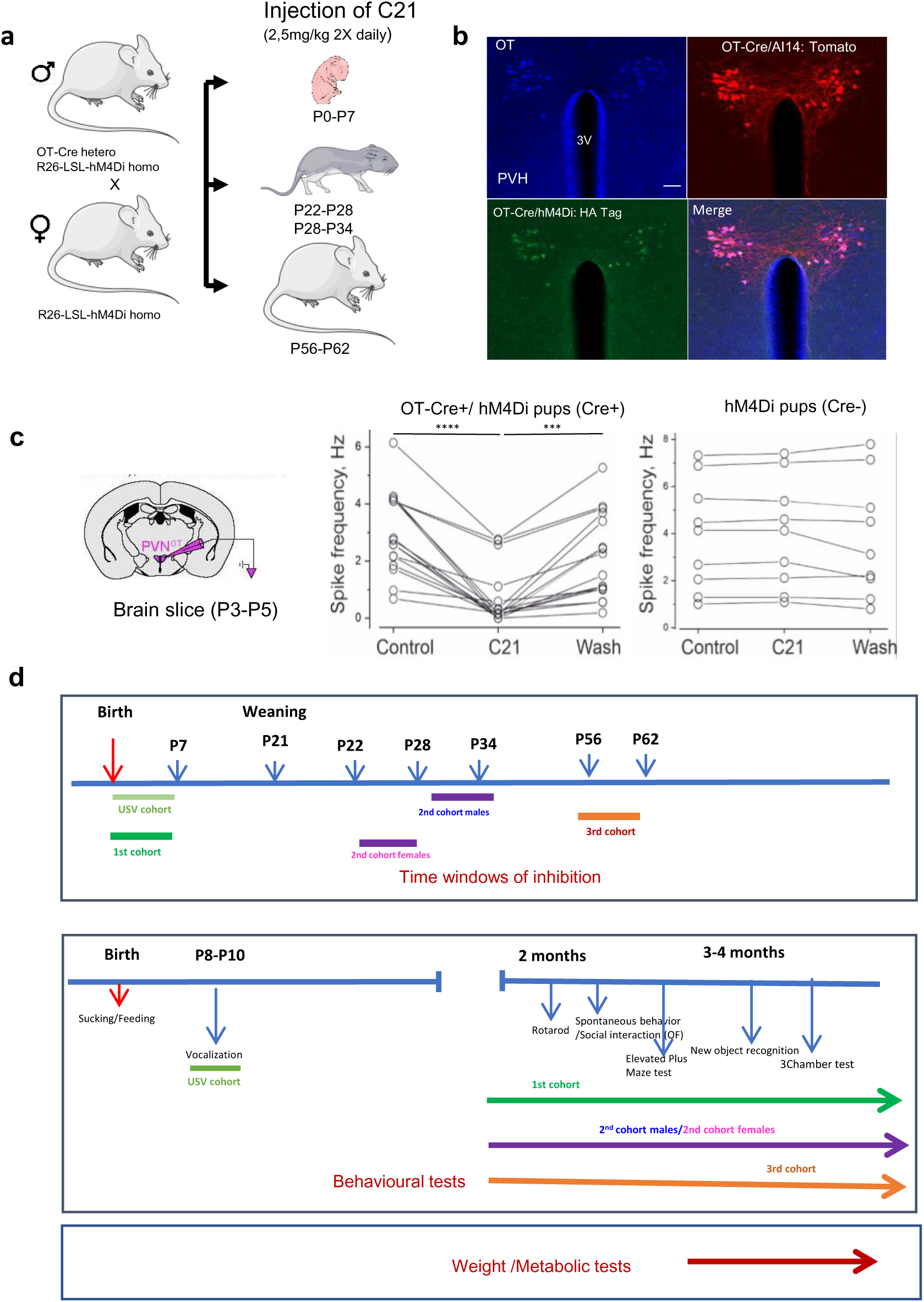
Chemogenetic inhibition of OT-expressing neurons at distinct postnatal stages and overall experimental design. (a) Breeding strategy and cohort generation producing Ot-Cre::hM4Di-HA (Cre+) and hM4Di-HA (Cre−) mice treated with the DREADD agonist C21 at defined developmental stages. (b) Immunohistochemical analysis of the paraventricular hypothalamus (PVH) in Ot-Cre::Ai14::hM4Di-HA mice showing oxytocin (OT, blue), tdTomato expression driven by Ot-Cre (red), and hM4Di-HA expression (green). Co-localization confirms selective expression of hM4Di-HA in OT-expressing neurons. Scale bar: 100 µm. (c) Whole-cell current-clamp recordings from tdTomato- and hM4Di-expressing OT neurons in slices from P3–P5 pups. C21 (10 µM, 5 min) markedly reduced action potentials in neurons from Cre+ mice but not Cre− animals. Spike frequency (Hz) in PVH OT neurons: Cre+ (N = 3 animals, n = 16 cells; Control: 2.95 ± 0.3, C21: 0.71 ± 0.2, Wash: 2.15 ± 0.3) and Cre− (N = 3 animals, n = 9 cells; Control: 3.93 ± 0.3, C21: 3.98 ± 0.5, Wash: 3.78 ± 0.3). (d) Experimental timeline indicating the timing of OT neuron inhibition and subsequent behavioral and metabolic assessments (rotarod, open field, elevated plus maze, novel object recognition, three-chamber test [males only], and body composition). Statistical analysis in (c) was performed using repeated-measures one-way ANOVA followed by Tukey’s multiple comparisons test. \*\*\**P* < 0.001, \*\*\*\**P* < 0.0001.

### Chemogenetic inhibition of OT-expressing neurons in neonates impairs ultrasonic vocalizations (USVs) in infancy and behavior in adulthood

To assess early communication, ultrasonic vocalizations (USVs) were recorded at P8 following OT-expressing neurons inhibition during the first postnatal week (Fig. 1d). Cre+ pups emitted fewer USVs than Cre− littermates (Fig. 2b), indicating impaired pup–dam communication [44, 45].

**Figure 2.**
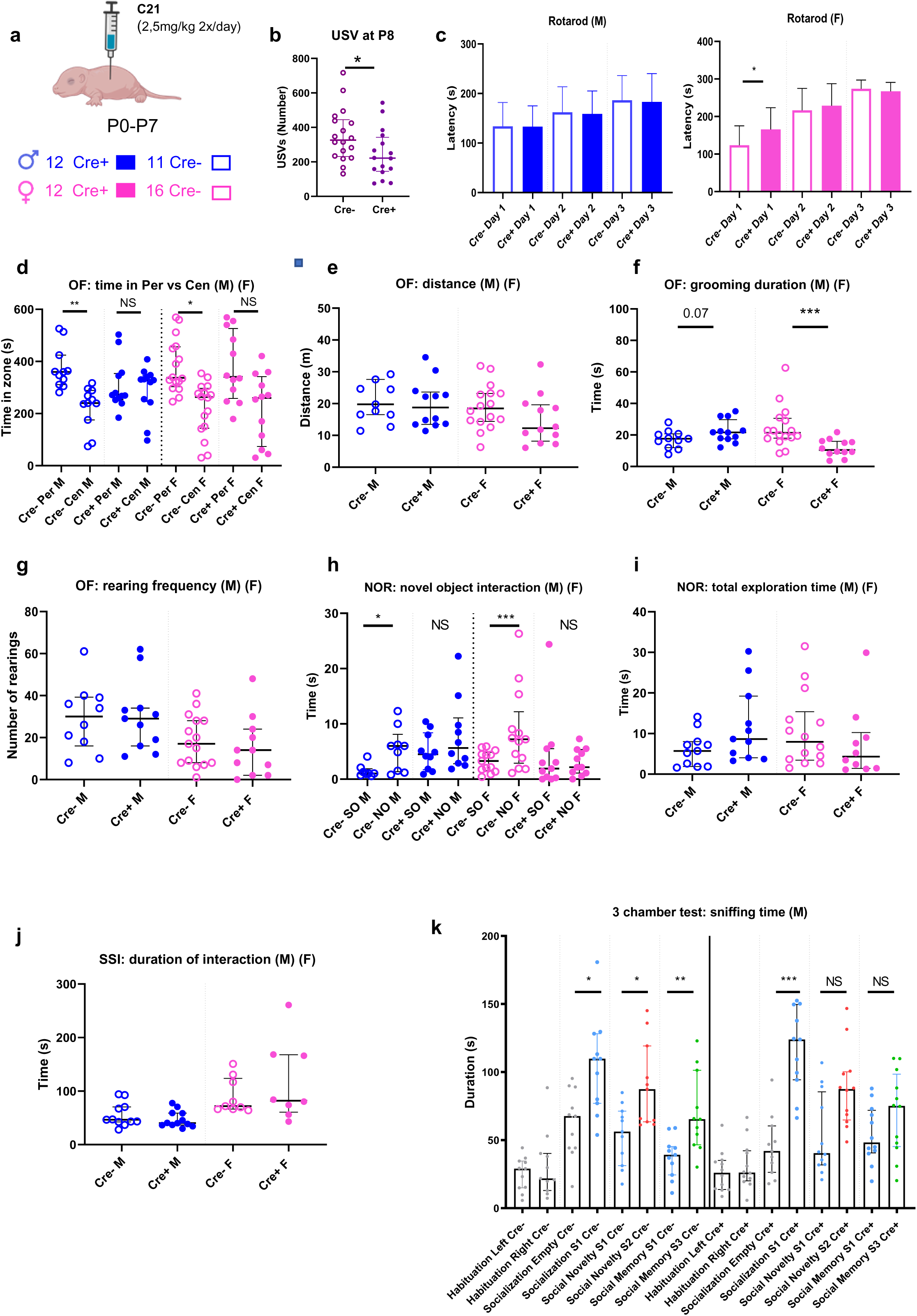
Behavioral effects of chemogenetic inhibition of OT-expressing neurons during infancy (P0–P7). (a) Mice received subcutaneous C21 injections during infancy (P0–P7). (b) Number of ultrasonic vocalizations (USVs) emitted by Cre+ and Cre− pups (males and females) at P8 following maternal separation. (c) Rotarod performance (latency to fall) in Cre+ and Cre− males and females across three consecutive days. (d–g) Open field test parameters including time in periphery versus center (d), total distance traveled (e), grooming duration (f), and rearing frequency (g). (h–i) Novel object recognition (NOR) test showing time interacting with familiar (SO) versus novel object (NO) (h) and total exploration time (i). (j) Spontaneous social interaction in the open field measured as time spent interacting with a same-sex juvenile conspecific. (k) Three-chamber test in males, showing time spent in each chamber during habituation and sniffing time directed toward each restrained mouse during socialization, social novelty, and social memory phases. Data are mean ± SEM for the rotarod test and median with quartiles for all other tests. Statistical analyses were performed as follows: Mann–Whitney tests (b, e, f, g, i, j), two-way ANOVA (genotype × sex) with Bonferroni post hoc test (c), Wilcoxon tests (d, h), or repeated-measures ANOVA (mixed-effects model) with Sidak’s multiple comparisons test (k). NS, not significant; \**P* < 0.05, \*\**P* < 0.01, \*\*\**P* < 0.001.

In a separated cohort subjected to the same neonatal inhibition, behavioral testing was performed in adulthood (Fig 2a). No major sensiromotor deficits were detected in the rotarod test over three days of testing, although Cre+ females showed a slight improvement on day 1 (Fig 2c). In the open field (OF) (Fig. 2d), male and female Cre− mice spent more time in the periphery (Per) than the centre (Cen) of the arena, whereas Cre+ mice of both sexes showed no compartment preference (Fig 2d). Locomotor activity (distance travelled in the OF) was unchanged (Fig. 2e), and elevated plus maze (EPM; Supplementary Fig. 1a) behavior was similar across groups. These findings suggest altered environmental processing rather than increased anxiety in Cre+ mice.

Self-grooming exhibited sex-dependent effects: increased in Cre+ males and decreased in Cre+ females (Fig. 2f). Self-grooming is considered a measure of repetitive behavior and involves the paraventricular hypothalamic nucleus (PVH) [46]. Rearing frequency was unaltered (Fig. 2g). In the novel object recognition (NOR) test, male and female Cre− mice prefererred the novel object, whereas Cre+ mice showed no preference despite similar total exploration time (Fig. 2h-i), indicating impaired object discrimination. Spontaneous social interactions (SSI) were unaffected (Fig. 2j).

In the three-chamber test (Fig. 2k), sociability was preserved in all groups. However, Cre+ males failed to show preference for social novelty and social memory was altered.

Together, these results demonstrate that neonatal inhibition of OT-expressing neurons reduces early communicative behavior and induces persistent deficits in environmental exploration and object recognition in both sexes, with male-specific impairments in social memory and social novelty preference. Anxiety-like behavior and basic sociability remain intact. In addition, genotype- and sex-dependent differences in self-grooming behavior were observed.

Altered social behavior is often associated with an imbalance between excitatory and inhibitory (E:I) activity in the hippocampus, as reported in a mouse model of autism involving neonatal oxytocin deficiency [25], in which neonatal OT treatment restored E:I balance [25]. We therefore performed whole-cell recordings in CA3 pyramidal neurons. No evidence of altered E:I balance was detected in juvenile Cre+ mice (Supplementary Fig. 2).

### Chemogenetic inhibition of OT-expressing neurons during puberty induces sex-specific behavioral alterations in adulthood

Male and female mice were treated with C21 during the pubertal period, accounting for sex-specific timing of puberty(Fig. 3a) [47, 48].

**Figure 3.**
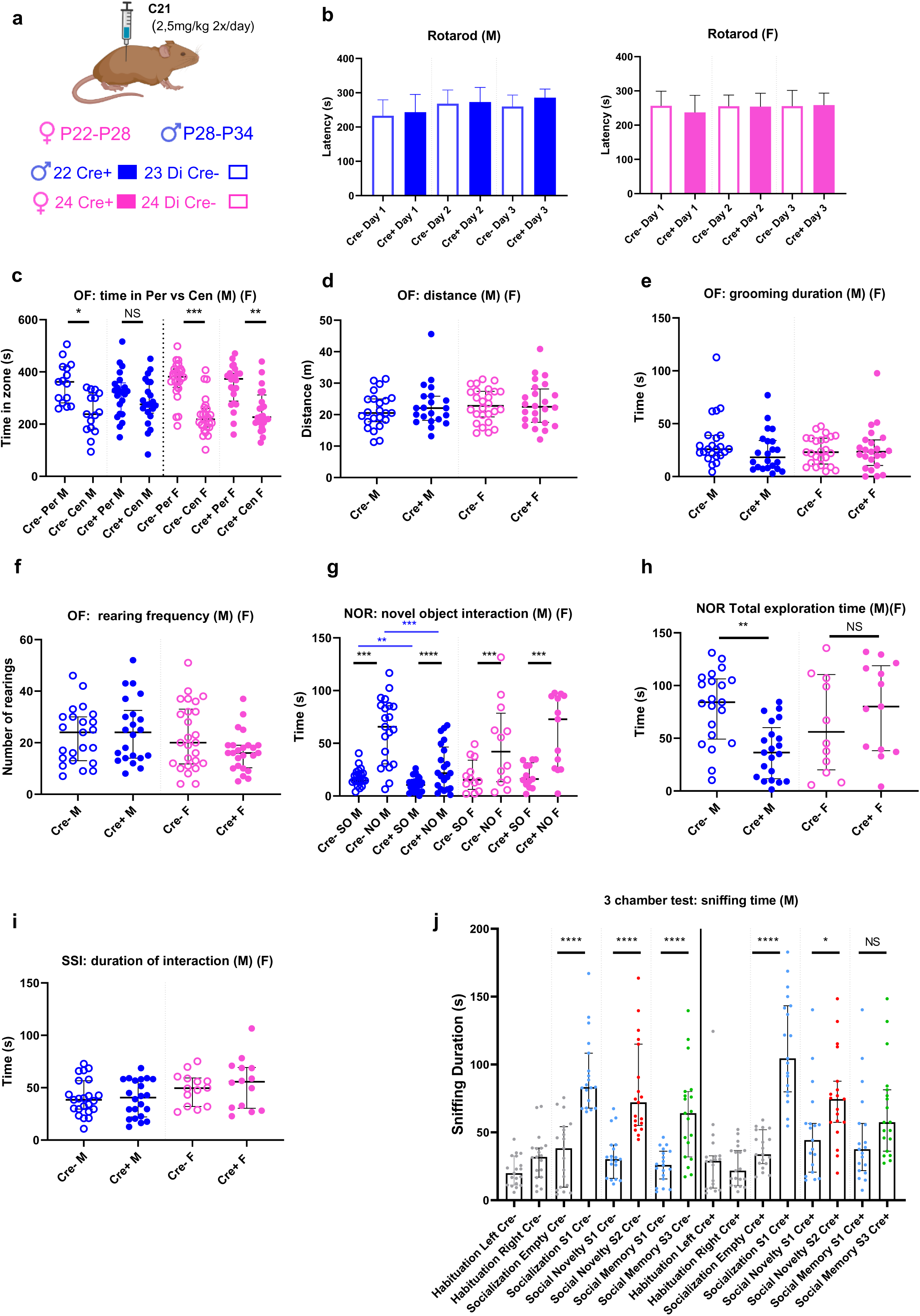
Behavioral effects of chemogenetic inhibition of OT-expressing neurons during puberty. (a) Cohorts received C21 during puberty (females: P22–P28; males: P28–P34). (b) Rotarod performance in Cre+ and Cre− males and females. (c–f) Open field test: time in periphery versus center (c), total distance travelled (d), grooming duration (e), and rearing frequency (f). (g–h) Novel object recognition test: time interacting with familiar versus novel object (g) and total exploration time (h). (i) Spontaneous social interaction in the open field. (j) Three-chamber test in males assessing sociability, social novelty, and short-term social memory. Data are mean ± SEM or median with quartiles. Statistical analyses were performed as follows: Mann–Whitney tests (b, d, e, f, h, i), two-way ANOVA (genotype × sex) with Bonferroni post hoc test (b), Wilcoxon tests (c, g), or repeated-measures ANOVA (mixed-effects model) with Sidak’s multiple comparisons test (j). NS, not significant; \**P* < 0.05, \*\**P* < 0.01, \*\*\**P* < 0.001.

No sensorimotor deficits were observed between Cre+ and Cre− males or females in the rotarod test (Fig. 3b). In the OF test, only Cre+ males lacked the typical preference for the periphery (Fig. 3c), while locomotor activity and EPM performance remained unchanged (Fig. 3d and Supplementary Fig. 1b), again suggesting altered exploration and/or environmental recognition in Cre+ mice rather than increased anxiety-like behavior.

Grooming duration (Fig. 3e) and rearing frequency (Fig. 3f) were unaffected. In the NOR test, all groups discriminated the novel object (Fig. 3f). However, Cre+ males exhibited reduced exploration duration (Fig. 3g) and total exploration time (Fig. 3h) for both familiar and novel objects. Spontaneous social interactions remained intact (Fig. 3i). but Cre+ males displayed impaired social memory and reduced preference for social novelty in the three-chamber test (Fig. 3j).

Thus, inhibition of OT-expressing neurons during female puberty had no detectable behavioral consequences in adulthood, whereas inhibition during male puberty selectively impaired exploration and social memory, resembling effects observed after neonatal inhibition.

### Chemogenetic inhibition of OT-expressing neurons in young adulthood produces selective behavioral effects in males

Male and female mice were treated with C21 between P56 and P62 (Fig. 4a). Behavioral testing was conducted at 2.5–4.5 months of age. No sensory-motor deficits were observed (Fig. 4b). All groups displayed normal anxiety-like behavior in the OF and EPM (Fig. 4c).

**Figure 4.**
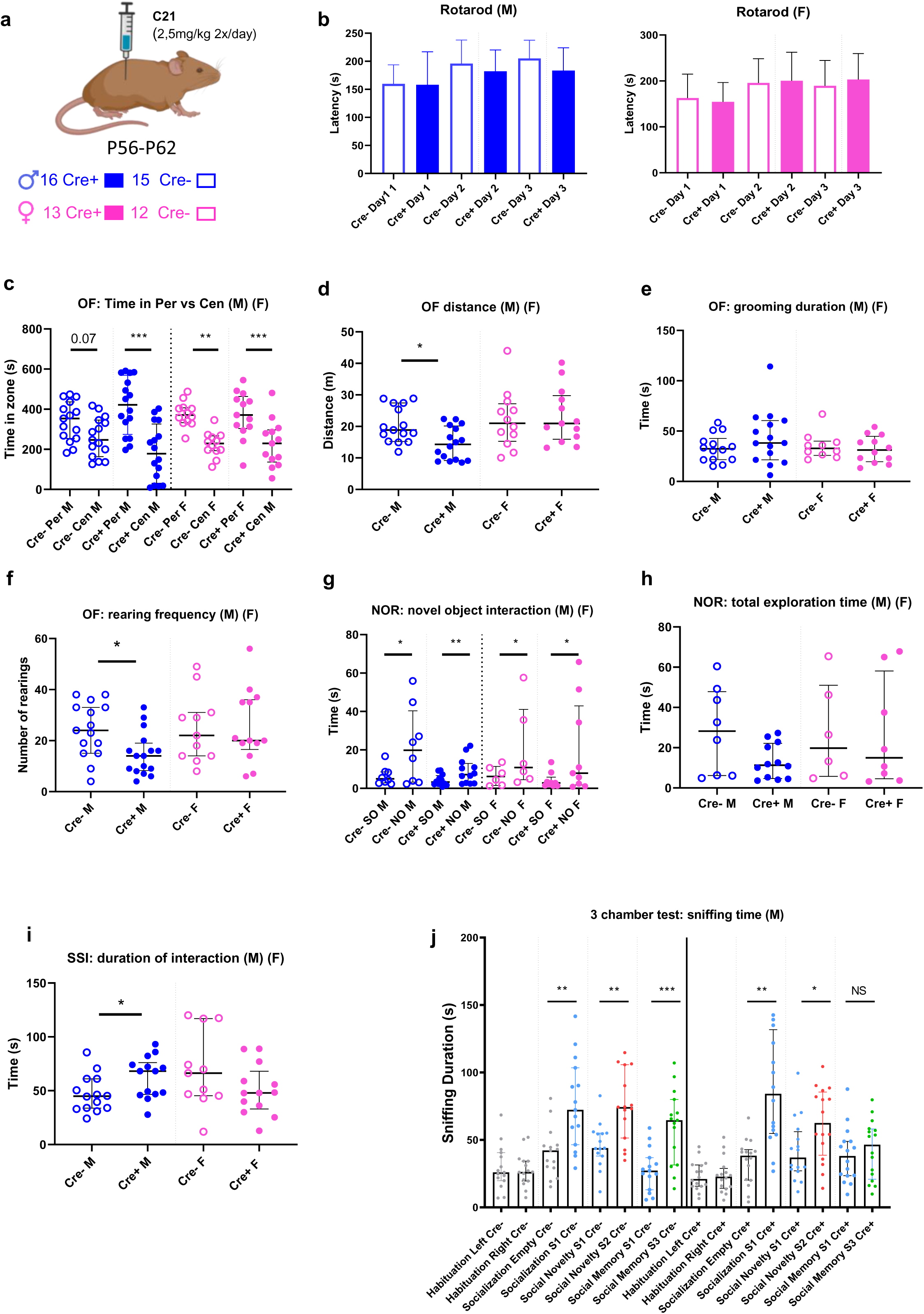
Behavioral effects of chemogenetic inhibition of OT-expressing neurons during adulthood (P56–P62). (a) Cohorts received C21 treatment during adulthood (P56–P62). (b) Rotarod performance in Cre+ and Cre− males and females. (c–f) Open field parameters: time in periphery versus center (c), total distance travelled (d), grooming duration (e), and rearing frequency (f). (g–h) Novel object recognition: time interacting with familiar versus novel object (g) and total exploration time (h). (i) Spontaneous social interaction in the open field. (j) Three-chamber test in males assessing sociability, social novelty, and social memory. Data are mean ± SEM or median with quartiles. Statistical analyses were performed as follows: Mann–Whitney tests (b, d, e, f, h, i), two-way ANOVA (genotype × sex) with Bonferroni post hoc test (b), Wilcoxon tests (c, g), or repeated-measures ANOVA (mixed-effects model) with Sidak’s multiple comparisons test (j). NS, not significant; \**P* < 0.05, \*\**P* < 0.01, \*\*\**P* < 0.001.

However, Cre+ males showed reduced locomotion (Fig. 4d) and fewer rearings (Fig. 4f), suggesting midly decreased activity. Objet recognition was unaffected (Fig. 4g–h). Cre+ males exhibite increased duration of spontaneous interactions with another mouse (Fig. 4i). yet showed impaired social memory in the three-chamber test (Fig. 4j).

These findings indicate that inhibition of OT-expressing neurons in young adult males induces subtle but long-lasting alterations in activity and social memory.

### Developmental timing of OT-expressing neurons inhibition affects body weight and adiposity in a sex-specific manner

Given the well-established anorexigenic role of oxytocin, we assessed body weight and adiposity in adulthood following inhibition at different developmental stages (Fig. 5a).

**Figure 5.**
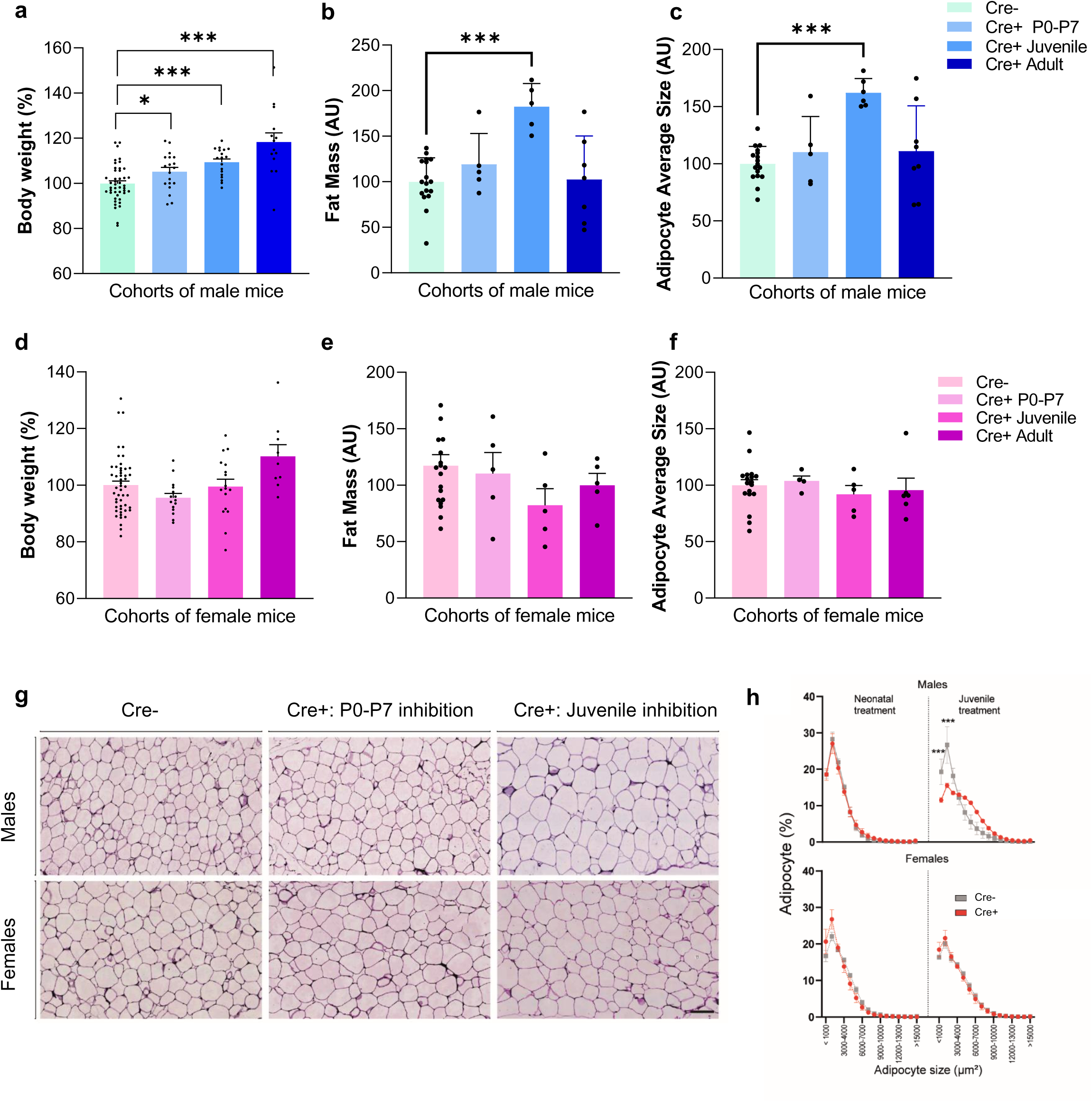
Effects of chemogenetic inhibition of OT-expressing neurons on body weight and adipose tissue across developmental stages. (a–c) Body weight (a), perigonadal adipose tissue mass (b), and mean adipocyte size (c) in 100-day-old males. (d–f) Corresponding measurements in females. Mice were treated with C21 during neonatal, juvenile, or adult stages. Values are normalized to Cre− controls. (g) Representative hematoxylin and eosin staining of perigonadal adipose tissue (scale bar: 100 µm). (h) Distribution of adipocyte size in male and female mice treated neonatally or during the juvenile period. Data are mean ± SEM. Statistical analyses were performed using Student’s t-test. \**P* < 0.05, \*\**P* < 0.01, \*\*\**P* < 0.001.

All Cre+ males exhibited increased body weight after inhibition during infancy (+9%), puberty (+17%), or young adulthood (+19%). However, increased fat mass (perigonadal adipose tissue, +82%) and adipocyte hypertrophy (+62%) were observed only following inhibition during puberty (Fig. 5b–c, g–h). In contrast, no metabolic alterations were detected in females across all conditions (Fig. 5d–f).

These results identify a male-specific sensitive window during puberty during which OT neuron inhibition promotes increased adiposity, whereas inhibition at other stages increases body weight without altering fat mass or adipocyte size.

### Chemogenetic inhibition of OT-expressing neurons in fetuses, from gestational day 18 to birth, delays parturition and impairs neonatal feeding

Previous studies have highlighted a critical role for OT in neonatal development, arising from both maternal release immediately prior to parturition [49] and endogenous production by the neonate from birth [50, 51]. To assess the role of OT signaling around birth, C21 (0.25 mg/ml) was administered in the drinking water of Cre− dams carrying mixed litters of Cre+ and Cre− fetuses (with paternal transmission of OT-Cre), from gestational day 18 (GD18) to birth (P0) (Fig. 6a). This strategy relies on the ability of C21 to selectively inhibit OT neurons in Cre+ fetuses/newborns.

**Figure 6.**
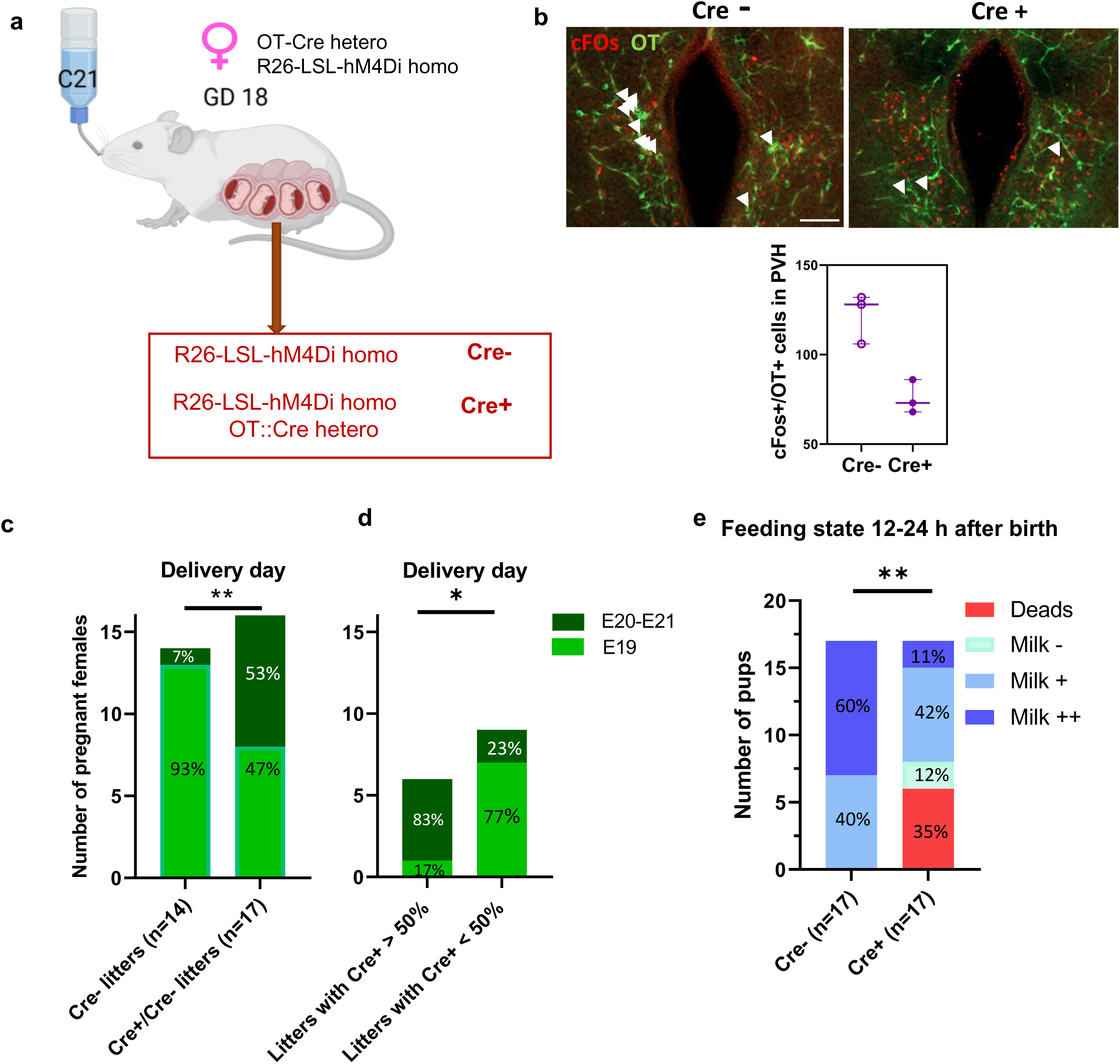
Prenatal chemogenetic inhibition of OT-expressing neurons and neonatal outcomes. (a) C21 was administered to dams via drinking water from gestational day 18 (GD18) until delivery (b) Immunohistochemical analysis of the paraventricular hypothalamus (PVH) in Cre− and Cre+ neonates 2 h after birth. cFos (red) and oxytocin (OT, green) labeling identifies activated OT neurons; quantification of co-localized cells indicates reduced OT neuron activation in Cre+ compared with Cre− neonates. Scale bar: 100 µm. (c) Delivery day of dams bearing litters composed exclusively with Cre− pups or mixed Cre+/Cre− pups. (d) Delivery day in dams with mixed litters containing >50% or <50% Cre+ pups. (e) Neonatal survival and feeding status at 12–24 h after birth, assessed by visual inspection of milk in the stomach. Statistical analyses were performed using the chi-square test. \**P* < 0.05, \*\**P* < 0.01, \*\*\**P* < 0.001.

Inhibition of OT neuronal activity in Cre+ neonates was confirmed by reduced c-Fos expression in PVH OT neurons after birth (Fig. 6b) that is typically observed in paraventricular hypothalamic (PVH) OT neurons within two hours after birth, following maternal care (F.M., unpublished data). Maternal C21 exposure markedly increased the incidence (53%) of delayed parturition (by 1–2 days) when litters contained Cre+ fetuses, compared with 7% in litters containing only Cre− mice (Fig. 6c). Notably, the delay positively correlated with the number of Cre+ fetuses (Fig. 6d), suggesting a contribution of fetal OT neurons to parturition timing.

Although we hypothesized that the fetal component of the placenta might express OT and thereby contribute to this process, no evidence of fetal OT expression was detected in the placenta at E18.5 (Supplementary Fig. 3).

Assessed of neonatal feeding revealed marked deficits in Cre+ pups 12–24 hours after birth: 35% died without milk in their stomachs, 12% were alive but had no milk, 42% contained only a small amount of milk, and 11% displayed a high milk content., compared with no mortality and normal feeding in Cre− pups (Fig. 6e).

These findings indicate that fetal/neonatal OT neuron activity contributes to both timely parturition and early neonatal feeding.

## DISCUSSION

The aim of this study was to determine whether discrete developmental windows exist during which OT-expressing neurons exert long-term effects on behavior and metabolism. To address this question, we performed chemogenetic inhibition of OT-expressing neurons across three key developmental periods (*i.e.,* infancy, the juvenile stage, and young adulthood), followed by longitudinal assessment of behavioral and metabolic outcomes in male and female mice. Across all time points, inhibition did not affect motor ability or anxiety-like behavior. In contrast, robust and specific alterations in social behavior were consistently observed. The severity and nature of these deficits varied as a function of both the developmental stage at which inhibition occurred and sex (see Table 1). The most pronounced behavioral impairements followed chemogenetic inhibition during the first postnatal week, affecting both males and females. Notably, social memory deficits were consistently detected in males, irrespective of the timing of inhibition. Body weight was affected in males at all stages of inhibition, whereas increases in fat mass and adipocyte size were restricted to inhibition during the juvenile period only. In addition, our finding unexpectedly reveal a role for fetal/newborn OT-producing neurons in the regulation of parturition timing and early feeding behavior.

**Table 1.** Behavioral effects of inhibition across developmental stages and sex.

| Test | Neonates<br>Males | Neonates<br>Females | Juveniles<br>Males | Juveniles<br>Females | Young<br>Adults<br>Males | Young<br>Adults<br>Females |
| --- | --- | --- | --- | --- | --- | --- |
| USVs | ↓ | ↓ | ND | ND | ND | ND |
| Rotarod | = | = | = | = | = | = |
| Open Field<br>(center time) | ↓ | ↓ | ↓ | = | = | = |
| Open Field<br>(distance) | = | = | = | = | ↓ | = |
| Open Field<br>(grooming) | ↑ | = | = | = | = | = |
| Open Field<br>(rearing) | = | = | = | = | ↓ | = |
| NOR<br>(discrimination<br>index) | ↓ | ↓ | = | = | = | = |
| NOR (total<br>exploration) | = | = | ↓ | = | = | = |
| Social interaction | = | = | = | = | ↑ | = |
| Three-chamber<br>test | = | ND* | = | ND | = | ND |

**Data are presented as direction of change compared to controls.**
↓ decreased; ↑ increased; = no significant difference; ND = not determined.
*ND*: test not performed or insufficient data.
NOR: Novel Object Recognition.
USVs: Ultrasonic Vocalizations.

Inhibition during the first postnatal week produced a more pronounced phenotype in both sexes. This induced reduced USVs, impaired novelty recognition, including difficulty distinguishing familiar from novel objects and our results suggested impaired recognition of the explored environment. In the three-chamber test, males maintained social interaction but failed to discriminate between familiar and novel conspecifics, and an impaired social memory. These phenotypes closely resemble those reported in adulthood in constitutive OT KO [29, 45, 52], OT-receptor KO [44, 53, 54], and CD38 KO mice [55, 56], CD38 being required for OT release. Together, these observations strongly suggest that the behavioural abnormalities observed in these transgenic models—characterized by transient inactivation of oxytocin-expressing neurons—are, at least in part, attributable to disrupted development of the oxytocin system, rather than to alterations in other factors produced by these neurons.

Importantly, early postnatal OT treatment has been shown to rescue adult social deficits in mouse models of ASD, particularly impairements in social memory [25, 27]. These long-lasting OT effects are thought to reflect normalization of critical neurodevelopmental processes within the hippocampus [25, 27], including the restoration of the developmental GABAergic shift, rebalancing of excitatory/inhibitory signaling, normalization of the number of somatostatin-expressing neurons [25], and regulating OT-receptor expression [57]. This early postnatal window coincides with the maturation and refinement of OT-related neuronal circuits. Our findings therefore provide strong evidence of a critical period during the first postnatal week during which OT-expressing neurons are essential for establishing lifelong social behaviors. Although inhibition during this period affected both sexes (with social memory only assessed in males), inhibition in juveniles or adults produced sex-dependent effects. This shift may be explained by the emergence of sexual dimorphism in OT release following the first postnatal week [58], driven by testosterone, which could underlie the sex-specific phenotype observed after infancy.

Inhibition during adolescence produced behavioral deficits exclusively in males, including altered exploration of the environment and objects, as well as impaired social memory. These findings may reflect sex-specific maturation of hypothalamic circuits and the establishment of functional dimorphism during puberty. Similarly, inhibition in young adults affected only males, leading to impaired social memory and modest reductions in locomotor activity and rearing in the open field. These results were unexpected, given that the OT system is considered mature at this age. Although the mechanisms remain unclear, one possible explanation is that transient inhibition of OT-expressing neurons in young adults induces epigenetic modifications that alter OT receptor expression or downstream signaling pathways, although this hypothesis requires further investigation.

Metabolic outcomes also depended on the timing of OT-expressing neurons inhibition. In males, inhibition during the juvenile period increased body weight, fat mass, and adipocyte size, whereas inhibition during infancy or young adulthood increased body weight without affecting adiposity. These findings suggest that OT regulates adult body weight across multiple life stages, while specifically influencing adipose tissue developemnt during the juvenile period through mechanisms that remain to be elucidated.

To investigate potential prenatal roles of OT neurons in fetuses, we inhibited OT neurons from gestational day 18.5 to birth via maternal administration of C21. This approach selectively targeted Cre+ fetuses and resulted in delayed parturition in dams carrying both Cre+ and Cre− fetuses, but not in dams carrying only Cre− fetuses. These findings show that fetal OT-expressing neurons contribute to the timing of delivery. While maternal OT is a well-established driver of myometrial contractions at parturition, studies in ewes and humans [59–62] have suggested a contributory role for fetal OT, potentially via fetal-to-maternal signalling and interactions with placental leucine aminopeptidase (P-LAP), which degrades OT [63]. In mice, however, mature OT production is generally thought to begin at birth [50, 64], and we did not detected OT expression in the placenta of E18.5 fetuses despite robust expression in the fetal brain. Further studies are therefore required to clarify the presence, source, and functional role of fetal OT in the mouse placenta prior to birth. In addition, 35% of Cre+ pups died on the day of birth due to failure to feed, and more than half had little or no milk in their stomachs. These findings indicate that neonatal inhibition of OT-expressing neurons disrupts suckling behavior and confirm our previous findings supporting a critical role for OT in the initiation of feeding in mouse neonates [51].

### Limitations

OT-expressing neurons are known to co-release additional signaling molecules, including neurotransmitters (glutamate), neuropeptides, and nitric oxide, which may contribute to the observed phenotypes. However, extensive prior literature detailed in the Introducion (and above) implicates OT itself as a key regulator of social behavior, and the ability of OT treatment to rescue social deficits with long-lasting effects in ASD models, strongly supports the interpretation that reduced OT signaling underlies the phenotypes observed here. Nonetheless, complementary approaches, such as temporarily contolled genetic inactivation of the OT gene, would help to further isolate the specific contribution of OT. Future studies should also incorporate additional behavioral and metabolic assays, and explore the molecular and circuit-level mechanisms underlying these effects in greater detail.

In conclusion, we identify distinct developmental windows during which OT-expressing neurons produce long-term effects on social behavior and metabolic regulation. These effects are both sex-dependent and stage-specific. Furthermore, our findings reveal previously unrecognized roles for fetal and neonatal OT-expressing neurons in the regulation of parturition timing and early feeding behavior. Together, these results refine our understanding of the developmental functions of the OT system and may inform the design of therapeutic strategies targeting these critical periods in neurodevelopmental disorders.

## AUTHOR CONTRIBUTIONS

Conceptualization: FM, SB; Methodology and Investigation: FS, CS, MSA, PYB, FO, JLG, YB, FB, EP, SGB, FM; Validation: FS, CS, MSA, PYB, SGB, FM; Resource: FM, SGB; Writing: FM; Visualization and Analysis: FS, PYB, MSA, CS, FO, JLG, YB, FM, SGB; Supervision: FM; Project administration: FM; Funding acquisition: FM and SGB. All the authors revised the manuscript.

## ACKNOWLEDGEMENTS

We thank the members of the animal facility of INMED laboratory and Marion Xolin for her technical help.

## Supplementary Figures

**Supplementary Figure 1:**
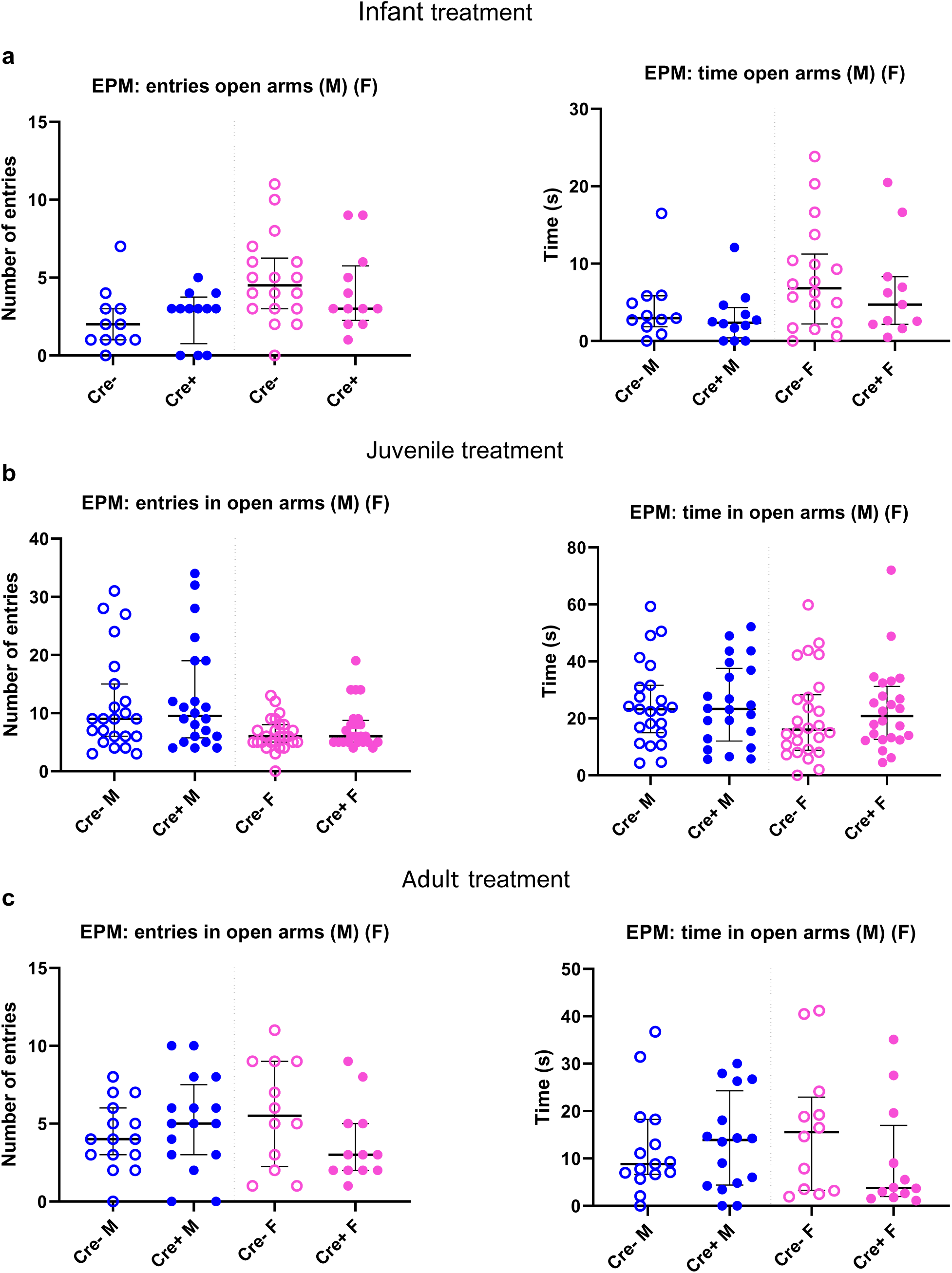
Elevated plus maze performance across male and female cohorts following chemogenetic inhibition of OT-expressing neurons. The number of entries into the open arms and the time spent in the open arms were quantified after inhibition at the neonatal stage (a), juvenile stage (b), and young adult stage (c). Data are presented as median and interquartile range. Statistical significance between groups was assessed using the Wilcoxon–Mann–Whitney test.

**Supplementary Figure 2:**
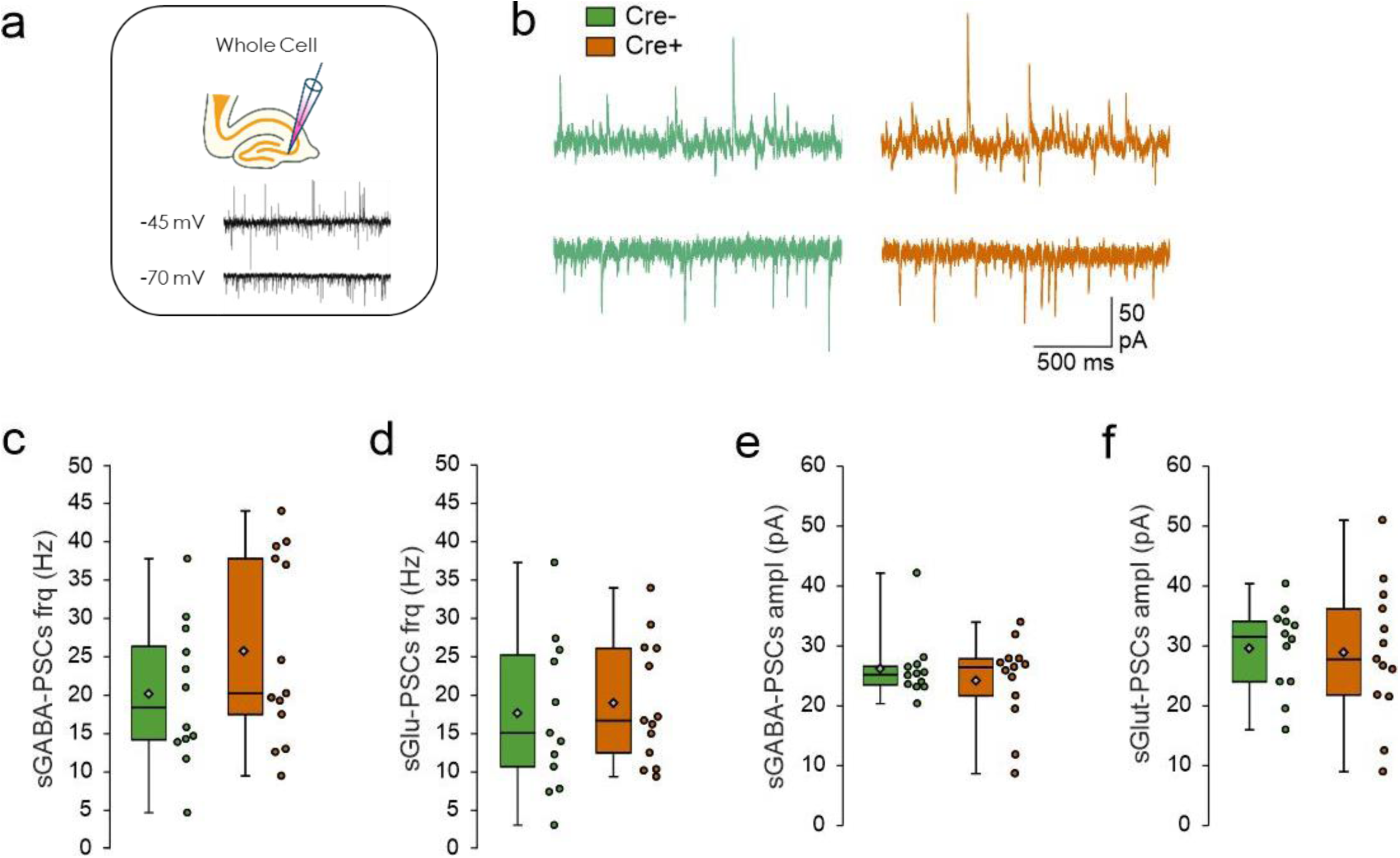
Spontaneous Glutamatergic and GABAergic synaptic activity of CA3 pyramidal neurons in the anterior hippocampus region in mice following chemoinhibition of OT-expressing neurons during infancy (Cre+) compared to controls (Cre-) mice. **a)** Schematic representation of the experimental protocol. Whole-cell recordings performed on CA3 pyramidal neurons were performed at a holding potential of −45 and -70 mV. The glutamatergic synaptic currents are inwards at -70 mV and the GABAergic synaptic currents are outwards at -45 mV. **b)** Examples of whole-cell recordings performed on CA3 pyramidal neurons at a holding potential of −45 and -70 mV at P50. **c-d)** Box plots of the frequency of GABA-sPSCs (**c**) and Glut-sPSCs (**d**). **e-f**) Box plots of the amplitude of GABA-sPSCs (**e**) and Glut-sPSCs (**f**). *Cre+* (N= 4, n=12) and Cre- (N=4, n=13) have been analyzed, with N: number of mice and n: number of recorded cells. Data represented in whisker-plots report the different values of recorded cells with mean ±SEM, with scattered plots showing individual data points: sGABA frq Cre- (20.1 ± 2.7); sGABA frq Cre+ (25.7± 3.3); sGlut frq Cre- (17.8 ± 2.8); sGlut frq Cre+ (18.9 ± 2.2); sGABA Ampl Cre- (25.7± 1.5); sGABA Ampl Cre+ (23.8 ± 2); sGlut Ampl Cre- (29.5± 2); sGlut Ampl Cre+ (28.8 ± 3.1). Statistical significance between groups was determined Mann Whitney test

**Supplementary Figure 3:**
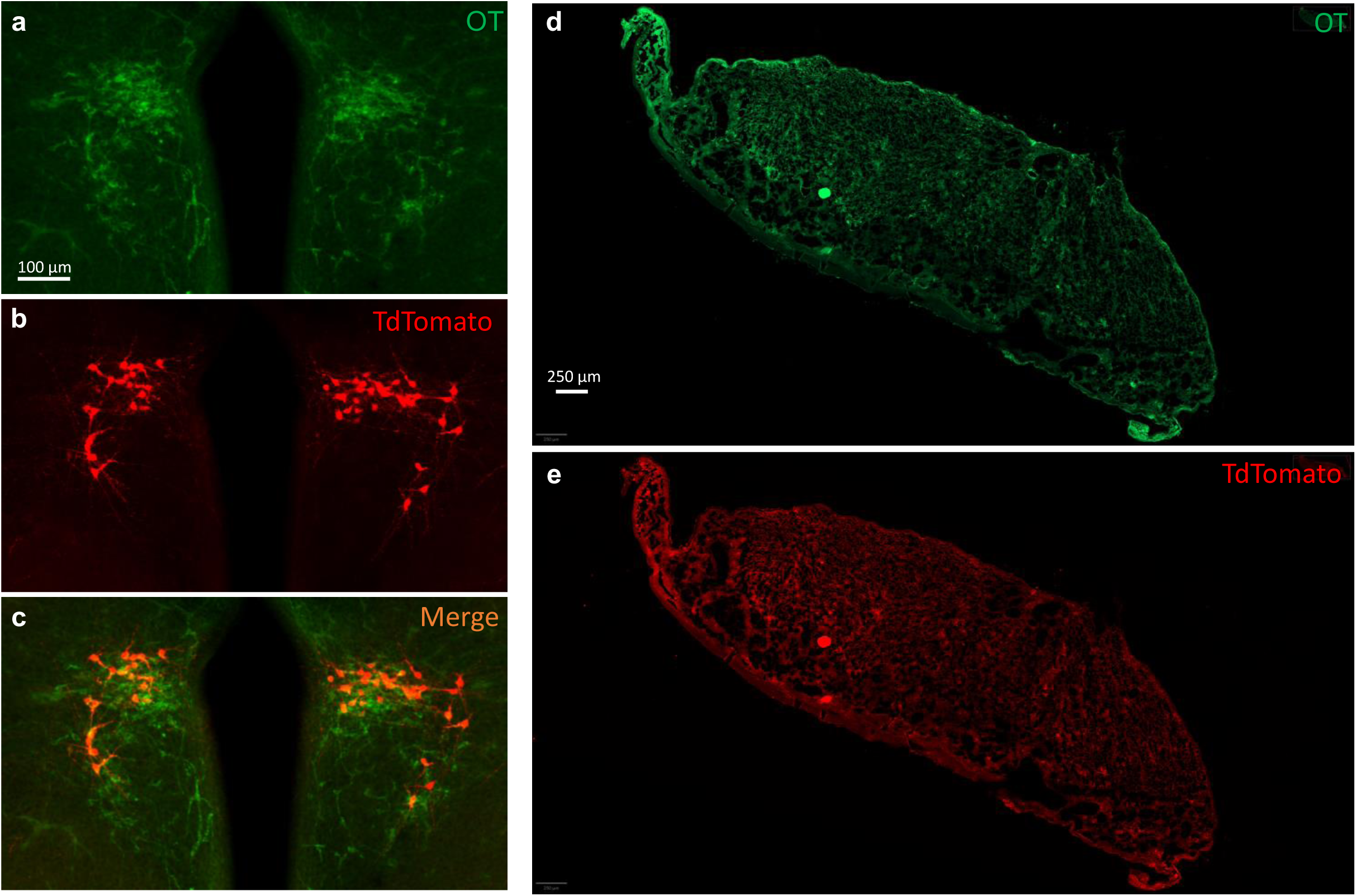
Fetal oxytocin expression in the brain and placenta of E18.5 OT-Cre::Ai14 fetuses. OT-Cre::Ai14 fetuses were generated by crossing an Ai14 (RCL-tdTomato) female with an OT-Cre male, allowing the visualization of OT expression via tdTomato fluorescence in fetal brain and placenta, but not in maternal tissues. OT expression, irrespective of maternal or fetal origin, was detected using an anti-OT antibody (green). (a, d) Immunohistochemistry using an anti-OT antibody in the brain (a) and placenta (d) of E18.5 fetuses. (b, e) Immunohistochemistry using an anti-tdTomato antibody in the brain (b) and placenta (e) of E18.5 fetuses. (c) Merged image in the brain.

## SUPPLEMENTARY INFORMATION

### SUPPLEMENTARY MATERIALS AND METHODS

#### Ultrasonic Vocalisations

Early social communication was tested by analyzing isolation-induced ultrasonic vocalizations (USVs) in P8 pups of both sexes{Bosque Ortiz, 2022 #5979}. Briefly, the mother and litter were left together for 30 min to habituate to the testing room, then P8 pups were all isolated from the mother and gently transferred to a new cage on a heating pad (37°C). After 5 min of separation, each pup was transferred in an isolation box located inside an anechoic box (54 x 57 x 41 cm; Couldbourn instruments, PA, USA). USVs were recorded for 300 s by an ultrasonic microphone (Avisoft UltraSoundGate condenser microphone capsule CM16/CMPA, Avisoft bioacustic, Germany) sensitive to frequencies of 10-250 kHz. Recordings were done using Avisoft recorder software (version 4.2) with a sampling rate of 250 kHz in 16-bit format. Data were analyzed for the total number of calls using Avisoft SASLab software.

##### Rotarod

Mice were challenged to walk on a rotating rod that increases in speed over a predetermined period of time (for 5 min) until the animal falls. The latency to fall from the rod is determined and taken as a measure of motor function. The test is performed over three successive days. The daily trial consists of three measures spaced by 5 minutes, a mean of the three measures is considered for each day.

##### Elevated plus maze

The device consists of a labyrinth of 4 arms of 5 cm width located 80 cm above the ground. Two opposite arms are open (without wall) while the other two arms are closed by side walls. The light intensity was adjusted to 20 Lux on the open arms. Mice were initially placed on the central platform and left free to explore the cross-shaped labyrinth for 5 minutes. The maze was cleaned and wiped with H_2_O and with 70% ethanol between each mouse. Animal movement was video-tracked using Ethovision software 11.5 (Noldus). Time spent in open and closed arms, the number of entries in open arms, as well as the distance covered, are directly measured by the software.

##### Open field

The open field test was performed in a 40 x 40 cm square arena with an indirect illumination of 60 lux. Mouse movement was video-tracked using Ethovision software 11.5 (Noldus) for 10 minutes. Total distance traveled and time in center (exclusion of a 5 cm border arena) are directly measured by the software. Grooming (time and events) and rearing were manually counted in live using manual functions of the software, by an experimented behaviourist. The open-field arena was cleaned and wiped with H_2_O and with 70% ethanol between each mouse.

##### Spontaneous Social Interaction in an open field arena

Animals were first habituated to the apparatus for 30 min. Social interaction was measured using pairs of mice from different housing cages, and having the same genotype, the same gender, and approximately the same body weight. Each pair was placed in the open field arena for 15 min during which different behavioural parameters reflecting social interaction between the two mice were recorded (following and sniffing). A social interaction index was defined as the percentage of counts and duration of social events over the total social and individual events.

##### New object recognition

First, two identical objects are placed in the arena, and then one of these objects is replaced by a novel object. Normal mice are expected to spend more time exploring the novel object than the familiar one. The arena used for the novel object recognition test was the same used for the open-field test. The arena was cleaned and wiped with 70% ethanol between each mouse. Two identical objects (50 ml orange corning tube) were placed in the opposite corners of the arena, 10 cm from the side walls. The tested mouse was placed at the opposite side of the arena and allowed to explore the arena for 10 min. After 1h, one object was randomly replaced with another novel object, which was of similar size but differ in the shape and color with the previous object (white and blue lego bricks), the other object (same object) was kept. Then, the same mouse was placed in the arena and allowed to explore the two objects (a new and an “old” familiar object) for 10 min. The movement of the mice was video-tracked with Ethovision 11.5 software. Time of exploration of both objects (nose located in a 2 cm area around object) was automatically measured by the software. The traveled distance to reach the novel object or the same object (old object) was measured.

##### Three-chamber social preference test

The three-chamber apparatus consisted of a Plexiglas box (50x25 cm) with removable floor and partitions dividing the box into three chambers with 5-cm openings between chambers. The task was carried out in four trials. The three-chambers apparatus was cleaned and wiped with 70% ethanol between each trial and each three-chamber test experiments. The mouse was video-tracked using Ethovision software. In a 5 minutes habituation trial, the tested mouse was placed in the central part of the 3-chamber unit and freely explore the two empty wire cages located in the left and right chambers (duration in each chamber). In the second 10 minutes socialization trial, a “stranger” wild-type mouse (same sex and same age, S1) is placed randomly in one of the two wire cages. The second wire cage remains empty. Social recognition is analysed during the third 10 minutes trial, in which a second “stranger” mouse (S2) is placed in the second wire cage. The S1 mouse is still under the first wire cage and corresponds to the familiar mouse. Thus, the tested mouse will have the choice between a familiar mouse (S1) and a new stranger mouse (S2). After a 30 minutes delay, the M2 mouse is replaced by another “stranger” mouse (S3) to test social memory. The measure of the real social contact is represented by the time spent in nose-to-nose interactions with the mouse. This test was performed using grouped-house mice.

#### Metabolic phenotyping

Animals were weighed at postnatal day 100 (P100) using an analytical balance. Perigonadal white adipose tissue was carefully dissected and weighed immediately after sacrifice using an analytical balance. Samples were fixed in 4% paraformaldehyde for 24 h at 4 °C and then dehydrated and paraffin-embedded using an automated tissue processor (Shandon Citadel™, Thermo Fisher Scientific). The dehydration sequence consisted of the following successive baths: 70% ethanol (1 h), 95% ethanol (1 h), 95% ethanol (1 h 30), absolute ethanol (1 h 30), absolute ethanol (2 h × 2), xylene (1 h), xylene (1 h 30), and paraffin (2 h 30 × 2). Paraffin used was Surgipath Paraplast (ref. 39601006), and xylene was Xylenes 98% pure (Acros Organics™, ref. 383930050).

Paraffin blocks were sectioned at 5 µm thickness using a Leica RM2245 microtome. Sections were collected on glass slides with a drop of water, stretched on a warm plate below the paraffin melting point (≤ 56 °C), and air-dried overnight at 35 °C. Slides were deparaffinized, rinsed in running water for 5 min, stained with Harris hematoxylin (Papanicolaou Harris hematoxylin, Bio-Optica, ref. 05-12011) for 3 min, rinsed again for 5 min, and counterstained with eosin Y 1% (1 g eosin Y, Acros Organics, ref. 152881000, in 100 mL Milli-Q water) for 3 min. Sections were dehydrated, cleared, and mounted with Eukitt. Slides were digitized using a Zeiss Axio Scan.Z1 automated slide scanner (Zeiss, Germany). Adipocyte morphology was quantified using ImageJ software (NIH, USA) with the home-made plugin to determine mean adipocyte area and size distribution.

